# Quantification of Adeno-Associated Virus with Safe Nucleic Acid Dyes

**DOI:** 10.1101/2020.02.27.968636

**Authors:** Jian Xu, Steven H DeVries, Yongling Zhu

**Affiliations:** Department of Ophthalmology, Northwestern University Feinberg School of Medicine, Chicago, IL 60611, USA

## Abstract

Adeno-associated virus (AAV) is the most commonly used viral vector for both biological and gene therapeutic applications^1^. Although many methods have been developed to measure quantity attributes of AAV, they are often technically challenging and time consuming. Here we report a method to titer AAV with GelGreen® dye, a safe green fluorescence nucleic acid dye recently engineered by Biotium company (Fremont, CA). This method, hereinafter referred to as GelGreen method, provides a fast (~ 30 minutes) and reliable strategy for AAV titration. To validate GelGreen method, we measured genome titer of an AAV reference material AAV8RSM and compared our titration results with those determined by Reference Material Working Group (ARMWG). We showed that GelGreen results and capsid Elisa results are comparable to each other. We also showed that GelRed® dye, a red fluorescence dye from Biotium, can be used to directly “visualize” AAV genome titer on a conventional gel imager, presenting an especially direct approach to estimate viral quantity. In summary, we described a technique to titer AAV by using new generation of safe DNA dyes. This technique is simple, safe, reliable and cost-efficient. It has potential to be broadly applied for quantifying and normalizing AAV viral vectors.

## Introduction

AAV is a small single-stranded DNA virus belonging to parvovirus family^2^. Features such as low toxicity, high safety, long-term expression and efficient transduction of both dividing and non-dividing cells have made AAV the most frequently used viral vector in biological studies^3–5^. Over the last two decades, AAV vector has also emerged as the most common vehicle for gene therapy^1, 6^. Back in 1995, an AAV vehicle was first used to treat cystic fibrosis in a human patient^7^. In 2008, three clinical trials using AAV vectors to treat Leber’s congenital amaurosis were published ^8–10^. As of to date, more than two hundreds clinical trials involving AAV have been conducted, whereby AAV vectors have shown great promise in treating different human diseases.

Recombinant AAV (rAAV) can now be packaged and purified quite routinely in laboratories, but their titers can vary largely, depending on packaging and purification methods and scales of production. Therefore it is imperative to establish accurate titers of rAAVs to ensure appropriate dosing. Many analytical methods, designed to measure either the physical or infectious titer of rAAV, have been developed. Among these are, for example, the dot blot hybridization^11^, enzyme-linked immunosorbent assay (Elisa)^12, 13^, Electron microscopy (EM)^14^, qPCR^15–17^, optical density^18^, DNA dye binding assay^19^ SDS-PAGE gel assay^20, 21^, TCID_50_ (50% Tissue Culture Infective Dose)^22^, replication center assay (RCA)^23^ and infectious center assay (ICA) assays^24^. Apparently each method has its own advantages and limitations. In the last several years, qPCR method has emerged as one of the most popular choices among labs, mostly due to its high sensitivity and broad dynamic range. But the qPCR method, like many others, also presents drawbacks. For instance, qPCR method is rather labor intensive. It is highly sensitive to experimental conditions, making it susceptible to errors. Factors such as PCR primers, reagents, equipment and DNA standards etc., can all significantly influence the test results^25, 26^. Because of that, significant inter- and intra- laboratory variations were often reported^27^. To overcome some of these issues, AAV titration method based on droplet digital PCR (ddPCR) was developed^27^. ddPCR is an endpoint PCR approach with the capability of measuring absolute number of DNA targets. Unlike qPCR, ddPCR is independent of reference materials and is less sensitive to inhibitors of PCR reactions, making it more accurate for measuring AAV titers^28^. However, ddPCR titration method is not widely used. Perhaps the requirement for special instrument and relatively high labor intensity have limited its broad application.

rAAVs can also be tittered by measuring their DNA contents more directly. For example, Cell Biolabs (San Diego, CA) has designed a commercial AAV titration kit (QuickTiter™ AAV Quantitation kit) based on quantifiable binding of DNA dye (CyQuant GR) to rAAV genome. Similarly, picoGreen, another sensitive DNA dye, has also been used to measure AAV titer based on the same principle^19^. A major advantage of DNA dye based assays is that they can be completed within 2-3 hours, much shorter than many other methods. Also, DNA dye-based assays was reported to have much less intra- and inter-assay variability as compared to dot blot and qPCR methods^19^. Notably, both CyQuant GR dye and picoGreen are membrane permeable dyes belonging to cyanine dye family. While CyQuant GR is often used in cell proliferation assays^29^, picoGreen dye is often used for quantifying double stranded (ds) DNA^30, 31^.

In recent years, several safe nucleic acid dyes have been developed, such as Gelgreen® and Gelred® from Biotium (Fremont, CA, USA), SYBRsafe and SYBRgold from Thermo-Fisher Scientific (Waltham, MA, USA) and Diamond™ from Promega (Madison, WI, USA). These dyes are now widely available as more and more labs are choosing them to replace ethidium bromide to stain DNA and RNA in gels. Compared with CyQuant GR dye and picoGreen dyes, these new dyes are more affordable. Importantly, they are membrane impermeant, making them safer to use and more friendly to environment.

In an effort to develop a safe, simple and reliable method for measuring AAV concentrations, we wondered if we could take advantage of the newly developed safe nucleic acid dyes, such as GelRed® and GelGreen®. According to Biotium, both GelRed® and GelGreen® can readily detect 1 ng of DNA in gel, with some users being able to detect bands containing less than 0.1 ng DNA. If these claims are true, then GelRed® and GelGreen® should at least be capable of detecting 3 - 4 μl of AAV at titer of 1 × 10^11^ GC/ml, which contains ~1 ng DNA (GC stands for genome copy; equations are provided in the method section). This level of sensitivity should be sufficient for most AAV samples as standard laboratory protocols typically produce rAAVs with titers one to two logs higher than 1 × 10^11^ GC/ml.

Here we report a method to measure AAV titer with GelGreen® and GelRed®. This method is fast, safe, reliable and cost-efficient. It produced similar result as compared to capsid Elisa method^32^. We believe this method could to be broadly useful in quantifying and normalizing AAV vectors.

## Materials and Methods

### rAAV production and purification

rAAVs were produced in-house using triple transfection methods^33–36^. The plasmids used for transfections were as follows: 1) cis-plasmid containing a gene expression cassette flanked by AAV2 inverted terminal repeats (ITRs); 2) trans-plasmids containing the AAV2 rep gene and AAV2 capsid protein genes; 3) adenovirus helper plasmid pAdΔF6. rAAVs were purified by iodixanol gradient ultracentrifugation as previously described^16, 37^. rAAV serotype 8 Reference Standard Material (AAV8RSM)^38^ was purchased from American Type Culture Collection (ATCC # VR-1816).

### Cytation 3 Plate reader

Cytation 3 Multi-Mode plate reader from BioTek (Winooski, VT) was used for DNA binding assay. It was equipped with a 488 nm laser for excitation and a 528/20 filter for emission. Gelgreen® was chosen to stain DNA because its excitation and emission spectrums are similar to GFP and it can be readily detected by virtually any plate readers.

### Gel imager

We used a DNA gels imager (Gel Logic 200 Imaging System) from Kodac (Rochester, NY), combined with a UV light box, to visualize viral DNA stained with Gelred®. Digital images were acquired and analyzed by image J software as described^39^ to provide a semi-quantitative analysis.

### Data Analysis

Statistical analyses were conducted with Graphpad Prism software (San Diego, CA). Student’s t-test and One-way ANOVA with Tukey’s post hoc test was used for data comparisons. Differences were considered significant when p < 0.05. Data are shown as mean ± SD.

The limit of detection (LOD) is defined as the mean value of sample blanks plus 3 standard deviations (SD). The limit of quantification (LOQ) is defined as mean value of sample blanks plus 10 SD.

A plasmid DNA, initially constructed as a cis-plasmid for making rAAV was used in this study as DNA standard. Its concentration was measured by NanoDrop™ Spectrophotometers (ThermoFisher, Waltham, MA, USA). The amounts of DNA (ng) in viral samples, either lysed or unlysed, were determined by standard curves. We then calculated the amount of encapsided DNA as the difference between the values of lysed samples and un-lysed samples.

The following are equations for converting encapsided DNA (ng) to AAV titer (GC/ml):

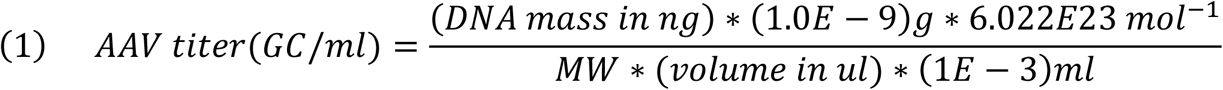

where,

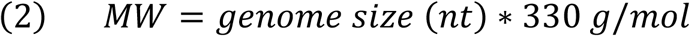

Note 1: 330 g/mol is the average mass of a single nucleotide (nt). Genome size of rAAV (ssDNA) is typically between 4000 to 5000 nt.

Note 2: If DNA sequence is available, MW of an AAV genome can be more precisely determined. For example, AAV8RSM was produced by pTR-UF-11 plasmid^40^. Based on its sequence, we calculated the MW of AAV8RSM’s genome to be 1 334 245 (g/mol). This number was used to compute titers of AAV8RSM in this report.

## Results

### Detection of DNA by Gelgreen®

To determine the detection limit and optimal concentration of Gelgreen dye, we carried out quantitative DNA binding assays using Cytation 3 plate reader. To set up the binding assay, we prepared several sets of DNA standards using a plasmid DNA. Each set of standard contains 12-point serial dilutions of DNA ranging from 0-50 ng. DNA standards were then transferred to 96-well plate containing GelGreen® diluted in phosphate-buffered saline (PBS) before fluorescence measurement.

Calibration plot of fluorescence intensity versus DNA is shown in Figure 1A (linear scale) and Figure 1B (logarithmic scale). Between 1/3000 to 1/100 000 dilution, GelGreen® readily responded to a wide range of DNA, showing linearity for DNA in the range of 0-50 ng, with all assays exhibiting acceptable correlation efficient (R^2^) of >99% (Figure 1A, 1B). However, at high concentrations of GelGreen® (1:500 and 1:1000), fluorescence signals no longer responded to DNA (Figure 1A, 1B). To view the effects of dye more directly, we re-plotted the data as fluorescence intensity vs. dye concentration in Figure 1C. This plot revealed a series of inverse bell-shaped dose-response curves for any given amount of DNA (0.78 - 50 ng), with their peak values all occurring at ~1/10 000 dye dilution and with sharp downslopes occurring after 1/3000. Thus in our assay, the optimal dye concentration for detecting DNA was 1/10 000, agreeing with manufacture’s recommendation.

**Figure 1.**
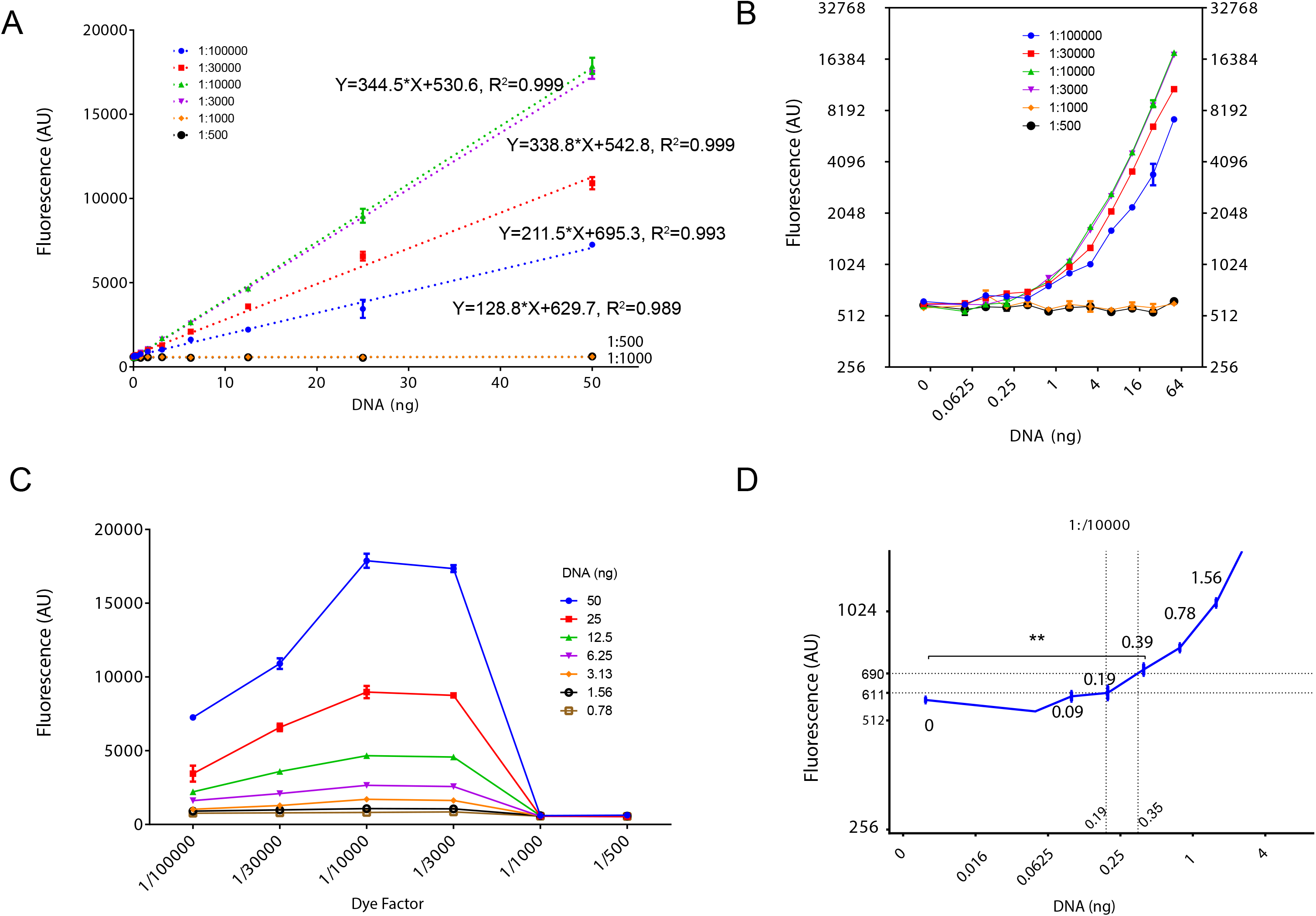
Detection of DNA by Gelgreen®. (A) Calibration plot of fluorescence intensity vs. DNA. Each line represents different dilutions of GelGreen®. Between 1/3000 to 1/100 000 dilution, GelGreen® showed linearity for DNA in the range of 0-50 ng. Results are averaged from triplicate wells. Bars indicate SD. (B) Plot of fluorescence intensity vs. DNA in logarithmic x-axis and y-axis. Each line represents different dilutions of GelGreen®. Results are averaged from triplicate wells. Bars indicate SD. (C) Effects of dye concentrations. Each line represents different amount of DNA. The results are averaged from triplicate samples. Bars indicate SD. (D) Limit of detection and quantification. GelGreen® was at 1/10 000. The plot shows an enlarged scale for DNA < 4 ng. Results are averaged from triplicate wells. Bars indicate SD. Limit of detection (LOD) (0.19 ng) and limit of quantification (LOQ) (0.35 ng) are marked with dashed lines. ** indicates statistically significant difference (t-test, p<0.01) compared to blanks (0 ng).

To assess the sensitivity of the DNA binding assay, we measured both the limit of detection (LOD) and the limit of quantification (LOQ) of GelGreen® at 1/10 000 dilution. (Figure 1D). With the standard deviation (SD) to be 10.60 and mean value to be 583 for sample blanks (n=3), we calculated the LOD and LOQ to be 611 and 690 respectively. Based on these values, we derived LOD to be 0.19 ng and LOQ to be and 0.35 ng from standard curve (marked with dashes lines in Figure 1D). In addition, using t-test we found that 0.39 ng DNA was the lowest amount to achieve statistical significance when compared to sample blanks (707.70 ± 26.03 vs 583 ± 10.60, n=3, p<0.01). Thus, Gelgreen® - based DNA binding assay is sensitive enough to measure as low as 0.2 - 0.4 ng DNA.

### Release of viral DNA by heating

AAV genome is encapsided. To measure it, one must first break apart viral capsids. Common methods for this purpose are proteinase K digestion^27^ and heat inactivation^18^. Often Proteinase K digestion is proceeded by DNAse I treatment to remove DNA contaminations^25, 27^. Heat inactivation is often performed around 70 °C in the presence of 0.05-0.1% SDS^18^. It usually takes one hour to perform these inactivation protocols.

In an effort to further shorten experimental time, we devised and tested two strategies for releasing viral DNA contents. The first strategy involves heating samples at high temperature of 95 °C, which can be done easily with standard PCR thermo-cycler. Meanwhile we also tested whether it is necessary to include SDS during heating process. In this experiment, we used an in-house produced rAAV sample packaged in capsid from serotype 2. We first diluted rAAV into 10 μl of PBS in PCR tubes then heated samples at 95 °C in thermo-cycler for various length of time. After heating and natural cooling, samples were transferred to 96-well plate containing 90 μl PBS and GelGreen® at 1/10 000 dilution for fluorescence measurement.

Results are shown in Figure 2A. We first inspected intrinsic dye signals (blanks) and compared that with signals of non-heated samples (see the blue line, 0 SDS). Sample blanks yielded signals of 515.7± 26.58 (n=3), whereas the non-heated sample exhibited slightly higher fluorescence (647.3 ± 16.80, n=3). The small difference here was thought to be caused by contamination of non-encapsided DNA in the AAV prep, which is quite common. Heating samples at 95°C caused a much larger increase of fluorescence signals, suggesting that contents of rAAV genome were released. Surprisingly, merely 5 minute of heating appeared to be sufficient, as longer heating (up to 20 minutes) produced no further increase of fluorescence. This observation implied that most of capsids, if not all, were already destroyed by heating after 5 minutes.

**Figure 2.**
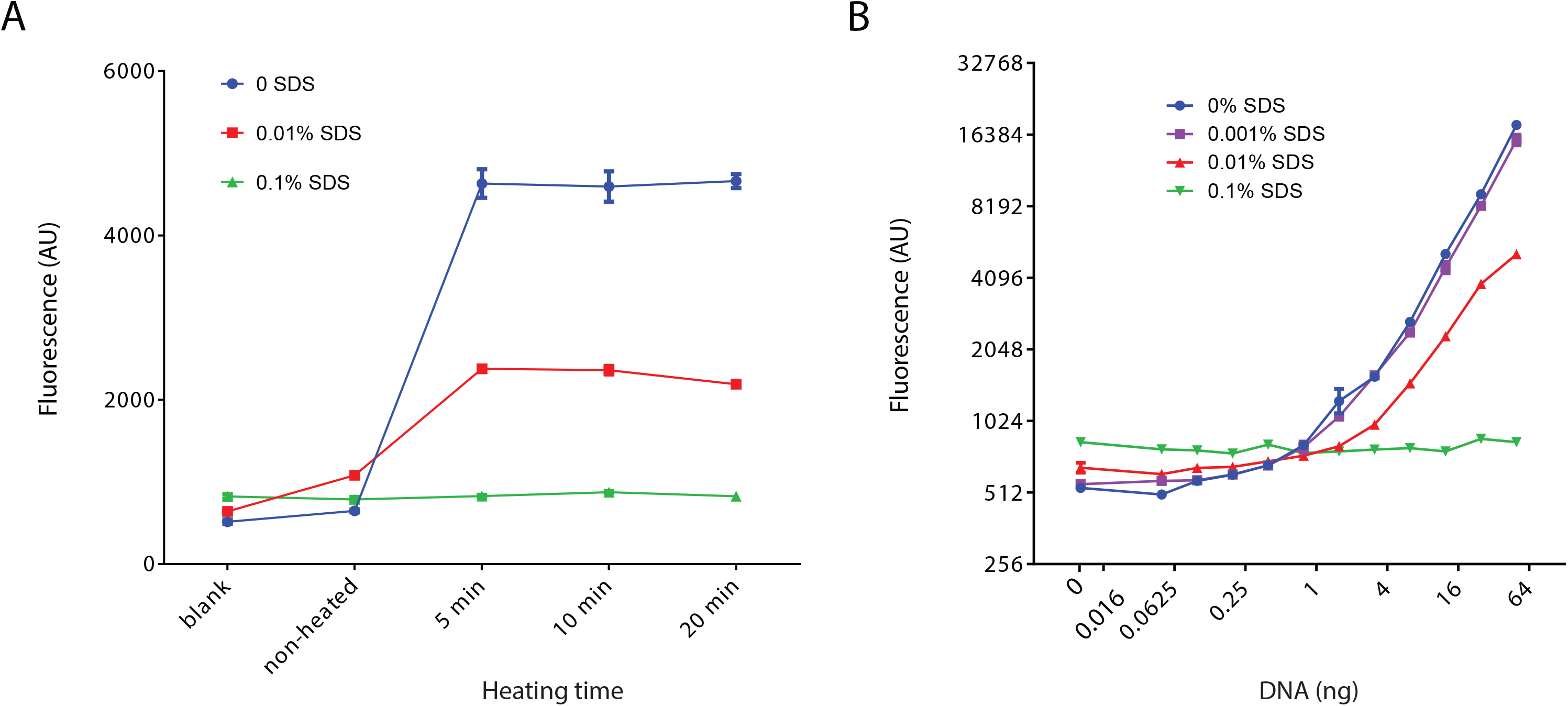
Release of viral DNA by heating at 95 °C. (A) Time course of fluorescence intensity during heating. Each line represents different concentration of SDS. Results are averaged from triplicate samples. Bars indicate SD. (B) Effects of SDS on DNA binding assay. Shown are series of curves of fluorescence vs DNA, with DNA ranging from 0-50 ng. Each curve represents different concentration of SDS. Results are averaged from triplicate wells. Bars indicate SD.

When samples (10 ul) were heated in the presence of 0.1% SDS, which yielded 0.01% SDS in final volume of 100 ul (red line), fluorescence signals again peaked at 5 minute. However these signals were significantly lower when compared to those without SDS (blue line). In an extreme case, when SDS was at final concentration of 0.1% (green line), the increase of fluorescence by heating was completely prevented (Figure 2A).

To investigate the effect of SDS more carefully, we prepared four sets of DNA standard containing different amount of SDS and performed DNA binding assays. Results are summarized in Figure 2B. While small amount of SDS (0.001%) had little effect as compared to control, 0.01% SDS already caused significantly inhibition. At 0.1% SDS, fluorescence responses completely vanished. Thus SDS must be kept low, otherwise will obstruct DNA binding assay. We advise that care should be taken when using GelGreen® to stain DNA in gels. Many GelGreen® users may not be aware that high amount of SDS (0.5% - 1%) is often included in 10X DNA loading dyes, which could have significant negative impact on detecting DNA bands by GelGreen®.

In conclusion, we found that 5 minutes of heating at 95 °C in PBS provided a simple and efficient way to release viral DNA. Meanwhile we also found that not only was SDS unnecessary but it was also detrimental for DNA detection by GelRreen®.

### Release of viral DNA by alkaline lysis

The second strategy we explored was the alkaline lysis. It is a very common molecular technique for protein and DNA denaturation, but it has rarely been used for lysing AAV particles^17^. We decided to test whether encapsided vrial DNA can be efficiently released by NaOH. Different amount of NaOH was tested for its lysing ability. The procedure is simple. We first treated viral samples (10 μl) with 2 μl of NaOH, we then added 2 μl of Tris buffer (PH 5.0) of two times the concentration of NaOH, to neutralize NaOH. Right after that we conducted DNA binding assay as described.

A dose-response curve of NaOH on fluorescence signals is shown in Figure 3A. Clearly, fluorescence signals were enhanced by NaOH, with large increases observed from samples treated with 15 mM to 125mM NaOH. It is likely that under these conditions viral particles were fully lysed. In comparison, 7.5 mM NaOH only caused a small fluorescence increase, suggesting that it only triggered a partial release. Based on the dose-response curve, we estimated the EC_50_ of NaOH treatment to be ~12 mM. Also noticeable was a small decline of fluorescence when NaOH was above 125 mM. We suspected that high dose of NaOH/Tris treatment may either cause DNA damage or weaken DNA/dye interactions. In light of this observation, we consider proper range of NaOH to be between 30 mM to 125 mM.

**Figure 3.**
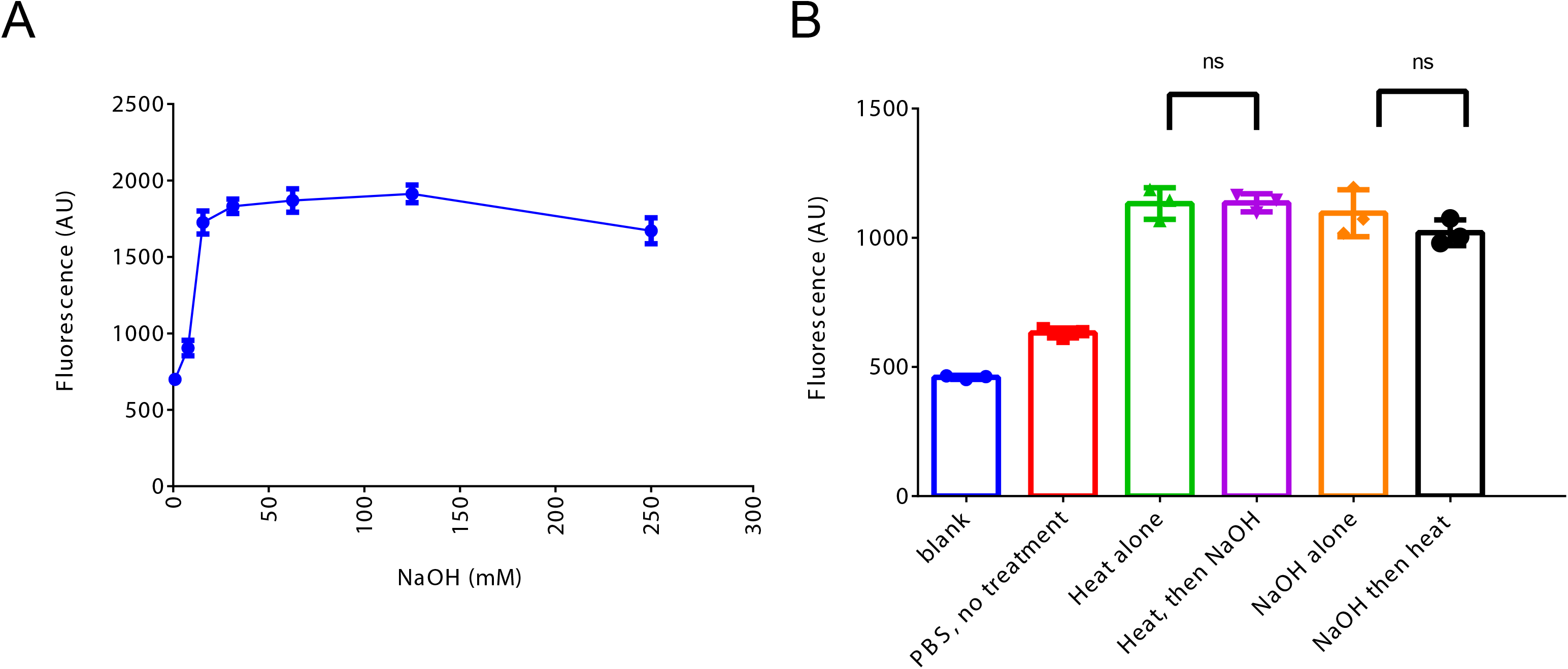
Release of viral DNA by alkaline lysis. (A) Fluorescence changes in response to NaOH treatment. NaOH ranges from 0 to 250 mM. Results are averaged from triplicate samples. Bars indicate SD. (B) Effects of combining alkaline lysis and heating procedures on viral lyisis. Results are averaged from triplicate samples. Bars indicate SD. ns = not significant.

Together we explored two methods to release viral DNA from capsids. Both methods can be done quickly and both appear to be fully effective. To be more assertive about their efficacies, we tested if combining two treatments could yield higher lysis than single treatment alone. Briefly, we either added NaOH (100 mM) to samples that have already been heated at 95 °C, or we subjected NaOH-treated samples to 95°C heating. As shown in Figure 3B, neither did NaOH (Purple) increase fluorescence to samples that have already been heated (Green), nor did heating (Black) enhance fluorescence of NaOH-treated samples (Orange). Together these results suggest that AAV were fully lysed by either of the two lysing methods alone, rendering the follow-up treatment nominal. In summary, heating method and alkaline method are both efficient and quick. Alkaline lysis is even easier to set up, giving itself a slight edge.

### Titration of AAV8RSM by GelGreen method

To validate GelGreen method for AAV titration, we measured titer of an AAV reference material (ARM) and compared our results to published results. The reference material was developed and characterized by ARMWG, for the purpose of normalizing titers of AAV vectors^38, 41^. Two ARMs, AAV2RSM and AAV8RSM, are available from ATCC (listed as ATCC-VR1616 and ATCC-VR1816, respectively). Their respective titers provided by ARMWG are 3.28 × 10^10^ GC/ml and 5.75 × 10^11^ GC/ml^38, 41^. We decided to use AAV8RSM for validation because its titer is higher and also because it has been extensively characterized. Many details were included in a series of publications^26, 32, 38, 42^, making it possible to compare our data with the literature values.

To measure the titer of AAV8RSM, we conducted three independent assays, with each assay performed in triplicates. For lysed samples, 1 μl of AAV8RSM was diluted in 10 ul PBS, followed by 2 μl of NaOH (500 mM) treatment for 2 minutes and then 2 ul of Tris (1M, PH 5.0) neutralization. For unlysed controls, 1 μl of AAV8RSM was simply diluted in PBS without addition of NaOH and Tris buffer. 12-point DNA standards ranging from 0 to 5.0 ng were also prepared. DNA binding assay was conducted as described before. Data from all three experiments was summarized in table 1.

**Table 1.** Titration of AAV8RSM (ATCC VR-1816)

| Intra assays |  |  |  |  |  |  |  |  |
| --- | --- | --- | --- | --- | --- | --- | --- | --- |
|  | Replicate<br>1 | Replicate<br>2 | Replicate<br>3 | Mean | SD | Low<br>95% CI | High<br>95% CI | CV% |
| Assay<br>#1 | 3.16E+11 | 3.46E+11 | 3.69E+11 | 3.44E+11 | 2.63E+10 | 2.78E+11 | 4.1E+11 | 7.7 |
| Assay<br>#2 | 4.71E+11 | 4.09E+11 | 4.89E+11 | 4.56E+11 | 4.18E+10 | 3.52E+11 | 5.6E+11 | 9.2 |
| Assay<br>#3 | 4.539E+11 | 3.41E+11 | 5.32+011 | 4.71E+11 | 1.1E+11 | 1.92E+11 | 7.5E+11 | 16.4 |
| Inter Assays |  |  |  |  |  |  |  |  |
|  | Mean | SD | Low 95%CI | High 95% CI | CV% |  |  |  |
|  | 4.23E+11 | 6.95E+10 | 2.51E+11 | 5.97E+11 | 16.3 |  |  |  |

Titer of the AAV8RSM was determined by GelGreen method. Three independent assays were conducted, with each assay being performed in triplicates. CV, coefficient of variation; CI, confidence interval.

An example to illustrate the analysis process is provided in Figure 4. In this experiment, unlysed samples exhibited fluorescence (527.5 ± 12.5, n=3) similar to sample blanks (516.8 ± 22.3, n=3), indicating that contamination of non-encapsided DNA was low in AAV8RSM. Based on the standard curve (Figure 4, inset), we estimated non-encapsided DNA to be 0.15 ng for each ul of AAV8RSM. Meanwhile lysed samples exhibited averaged fluorescence of 762.7± 27.42 (n=3), translating to 1.20 ng of DNA per ul of AAV8RSM. Therefore, encapsided DNA, calculated by subtracting values of unlysed samples from value of lysed samples, equals to 1.05 ng/ ul virus. Based on the provided equation, titer of AAV8RSM from this experiment was calculated to be 4.7×10^11 GC/ml.

**Figure 4.**
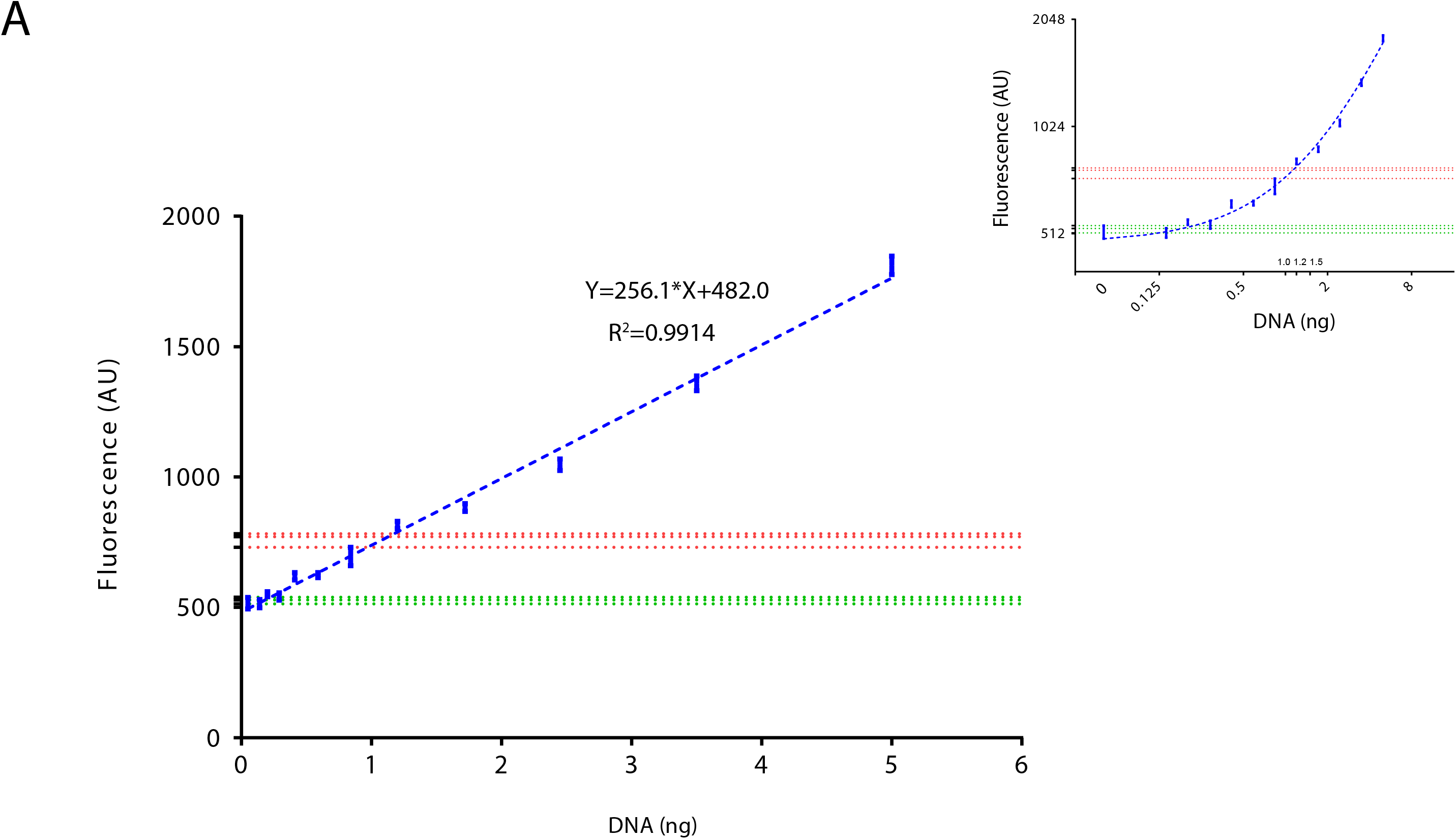
Titration of AAV8RSM by GelGreen method. (A) Calibration plot of standard DNA in linear-linear scale and in log-log scale (inset). The DNA standard was made by serial dilution from 5 ng with dilution effect of 0.7. Trend lines, linear regression equations and R^2^ are shown. Dash lines are for three untreated rAAV sample intersecting Y-axis at 514, 529 and 539 (green), and three NaOH-treated samples at Y-axis of 771, 732 and 781 (red) respectively.

In the same way we calculated titers of AAV8RSM for each independent assay (Table 1). At first glance, titers determined in each replicate, from all three experiments, fall into the range of 3-5 × 10^11^ GC/ml, similar to those obtained by Elisa method in previous studies^26, 32, 38, 42^. We will discuss this in more details in the discussion section. Intra-assay analysis was performed and it revealed low coefficients of variation (CV), with the highest being 16.4% and lowest being 7.7% (Table 1). Inter-assay analysis of three independent experiments showed mean titer of 4.23 × 10^11^, with low 95% confidence interval (CI) and high 95% CI to be 2.50 × 10^11^ and 5.97 × 10^11^ respectively, and with inter-assay CV to be 16.3%. Taken together, coefficients of variation of both inter-assay and intra-assay are quite low, indicating high repeatability and reproducibility of the GelGreen method (Table 1).

### Evaluation of the accuracy of GelGreen® - based AAV titration method

To further evaluate accuracy of the GelGreen method, we designed and performed a new experiment, in which we adopted the concept of amplification efficiency from qPCR analysis. The amplification efficiency of qPCR is calculated as E = −1+n^(−1/slope)^ where n is dilution factor and slope can be derived from linear regression of threshold cycle (Ct) vs. log of input DNA. In general, efficiency between 90% and 110% is acceptable.

In a similar way, we measured efficiency of GelGreen method using serial 2-fold dilution of a home-made rAAV sample. We used NaOH to lyse viral particles and conducted DNA quantification as before. DNA standard curve is shown in Figure 5A. As expected, fluorescence of serially diluted AAV samples progressively declined (Figure 5B). Similarly, DNA mass, which was converted from florescence based on standard curve, also took steps down during serial dilution (Figure 5C). We recognized that 1 μl of viral sample contained ~18 ng DNA. A simple calculation yielded a titer of 6.99, × 10^12^ for this sample (Figure 5C). After six rounds of two-fold dilution, total DNA was reduced to less than 0.3 ng and consequently became non-detectable (Figure 5 B-D). Meanwhile, linear relationships existed between Log_2_(DNA) and number of dilutions up to 6 (Figure 1D). Slopes from three independent serial dilutions were derived from the linear portion of the curves to attain a mean value of −0.96 ± 0.04. Accordingly, we calculated efficiency of GelGreen method to be 106 ± 11 %, which is very close to the theoretical 100% efficiency.

**Figure 5.**
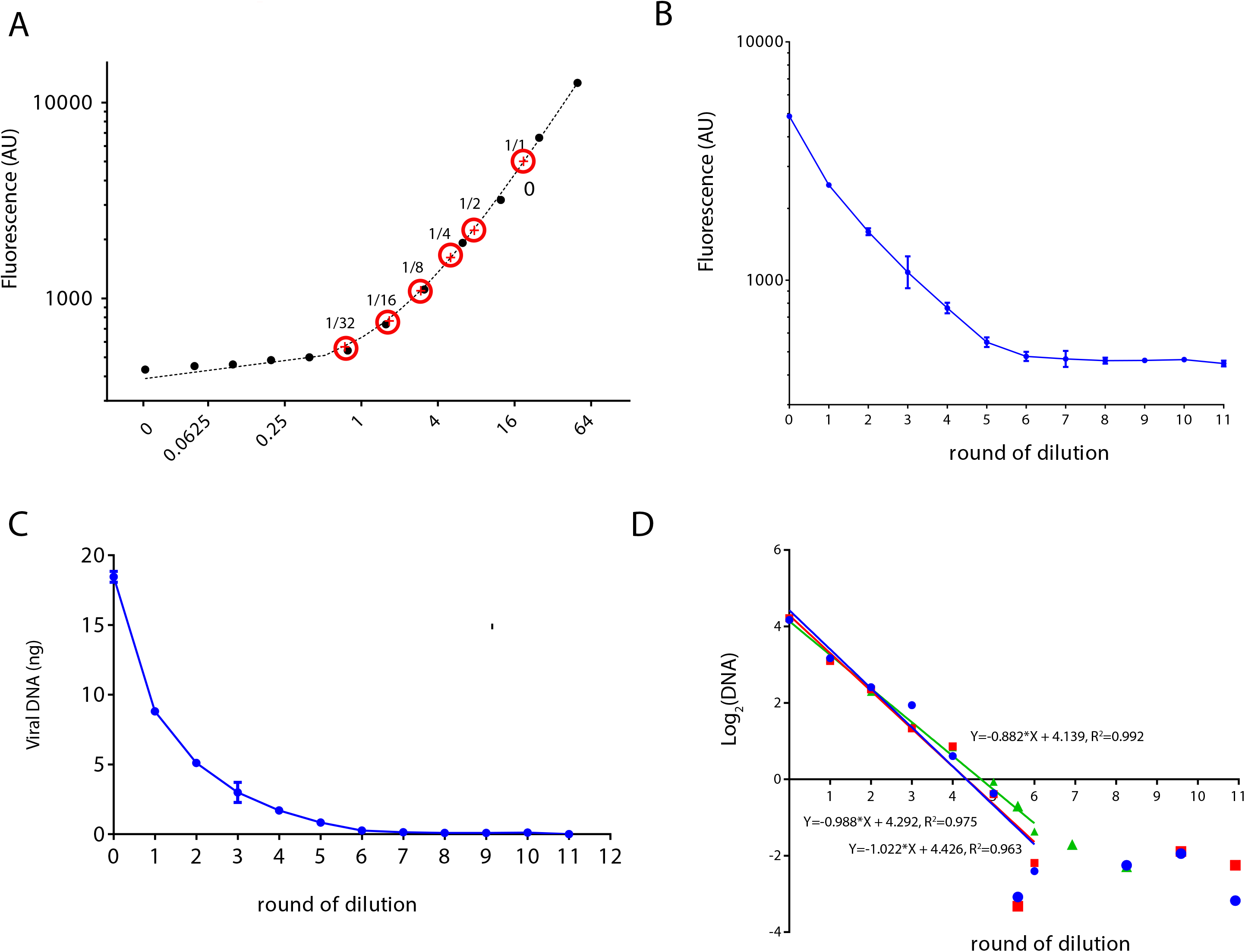
Evaluation of the accuracy of the GelGreen-based AAV titration method. (A) DNA standard curve (log-log plot). DNA standard was made by two-fold serial dilution, ranging from 0 to 50 ng. Red circles on the fitted line indicate fluorescence of AAV virus (averaged from triplicate samples) at different dilutions. (B) Fluorescence of serially diluted AAV virus. Results are averaged from triplicate samples. Bars indicate SD. (C) Plot of converted viral DNA mass vs. dilutions. Results are averaged from triplicate samples. Bars indicate SD. (D) Plot of Log2 of the DNA vs. dilutions. Each line represents one dilution series (n=3). Linear relationships exist when the dilution rounds were less than 6. Trend lines, linear regression equations and R^2^ are shown.

### Visualization of rAAV quantity with gel imager

Having measured AAV titer with plate reader, we wondered if we can even “visualize” AAV titer directly with a gel imager. We are equipped with a system intended for imaging ethidium bromide (EB) stained DNA. Thus we chose GelRed® in this experiment because its fluorescence properties are similar to that of EB. A pilot experiment found that 1/10 000 dilution of GelRed® is also the optimal dilution factor for DNA detection, like GelGreen®.

We selected a rAAV sample whose titer was about 4.5×10^12^. We made four-point serial dilution (2-fold) of virus in a PCR strip, with the starting tube containing 1 μl virus diluted in 10 μl of PBS. Samples were heated at 95 °C for 5 minutes in a PCR thermo-cycler. After heating, we pipetted 5 μl of 1/3,300 GelRed® into PCR strip to make final 1/10 000 GelRed® dilution. We also made DNA standard (0-50 ng) in a PCR strip. Viral samples and DNA standard were imaged simultaneously with gel imager (Figure 6A). By side-by-side comparison, it is quite easy to approximate that the amount of DNA in the starting tube is between 6.26 to 12.5 ng. To be more quantitative, we analyzed image file with image J software to generate a calibration plot. This plot shows linear relationship between fluorescence intensity and DNA up to 25 ng (Figure 6B). Based on the standard curve, we calculated the DNA content in the staring tube to be 11.9 ng. Similarly, viral sample’s DNA contents at each dilution were derived. The plot of Log_2_(DNA) vs dilution showed a linear relationship with a slope of −0.87, which yielded a 121% efficiency (Figure 6C). Thus, simple imaging method provided a quick, and fairly effective way to estimate AAV titers.

**Figure 6.**
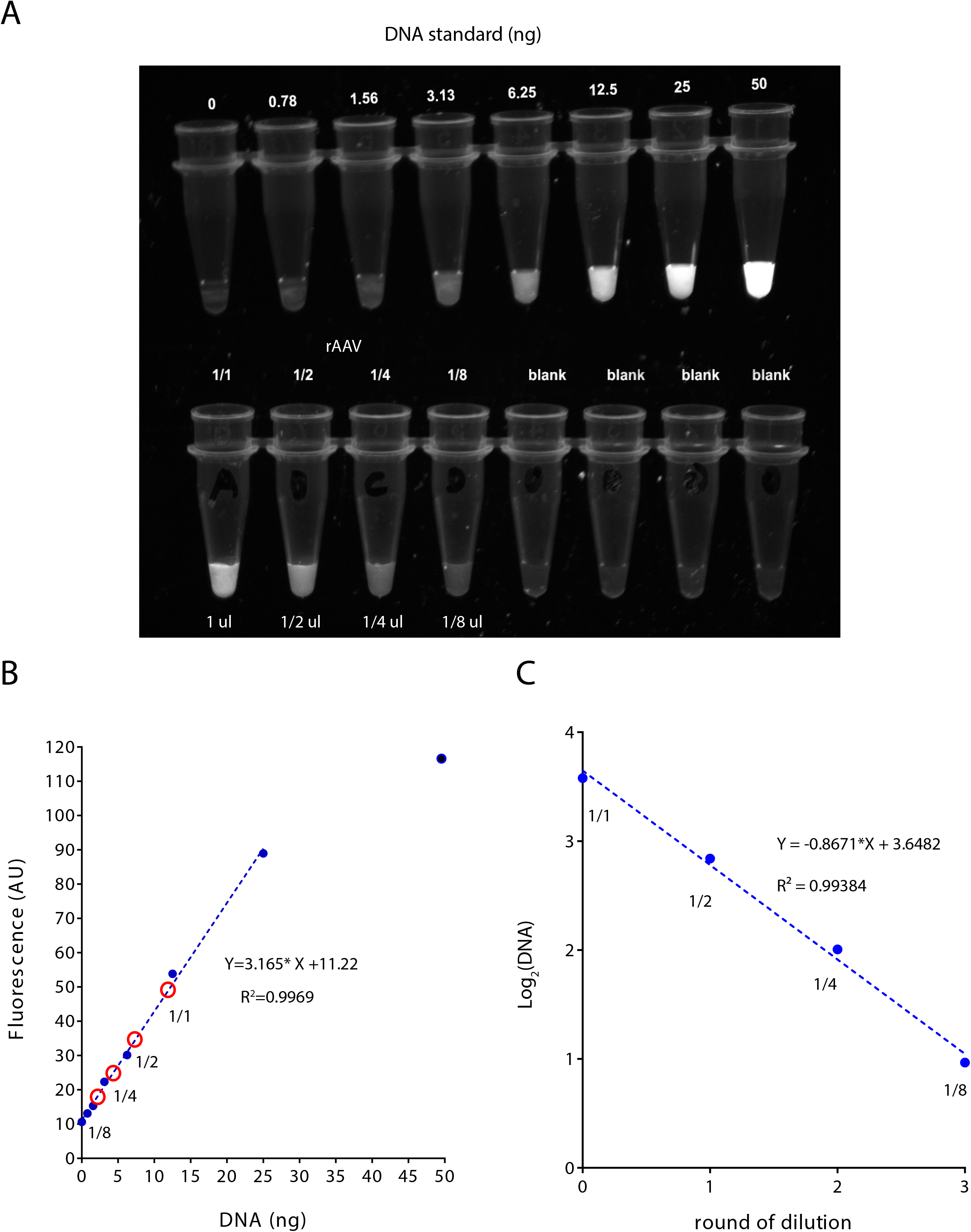
Direct visualization of quantities of rAAV using gel imager. (A) Images of DNA standard (top) and AAV samples (Bottom) stained by 1/10 000 GelRed®. DNA standard was made by two-fold serial dilution, starting at 50 ng. AAV samples consisted of 4-point twofold dilution series. (B) Calibration plot of DNA standard. GelRed® shows linearity for the range of 0-25 ng DNA. Trend lines, linear regression equations and R^2^ are shown. Red circles on the fitted line indicate fluorescence of serially diluted vial samples. (C) Linear approximation of Log2 DNA input vs. dilution. Calculated slope and R^2^ are shown.

## Discussion

Here we report a method to quantify AAV vectors based on binding of AAV’s DNA with a safe nucleic dye. This method offers several advantages. First, it is very fast. It allows determination of AAV titrations in about 30 minutes. Second, it is safe and cost-efficient. The Biotum dyes we used are membrane impermeant, making them safer to use and less hazardous to environment. It is also economical. Each experiment typically requires less than 1 μl of dye, costing only 20-30 cents. Most importantly, this method is consistent, as inter-assay and intra-assay variations are both small. We believe this is mainly due to the fact that DNA was measured directly without amplification. Skipping amplification steps makes the assay less sensitive to many factors that are crucial for enzyme-based reactions, such as PCR and capsid Elisa. The main disadvantage of the method is its low sensitivity, at least when compared to qPCR method and Elisa. Since LOD of GelGreen is ~ 0.3 ng, we estimate that the lowest AAV titer this method can detect is about 1.0 × 10^10^ gc/ml at the expense of 10 μl of viral sample.

AAV genome is single-stranded DNA (ssDNA) of approximately 4.7-kilobases (kb). It was flanked with two 145 nucleotide-long inverted terminal repeats (ITR) that actually form double stranded DNA (dsDNA). Thus in its natural form, AAV genome is made of both ssDNA and dsDNA. Although it has been suggested that following denaturation AAV genome anneals to form dsDNA^18^, the extent to which dsDNA is converted from ssDNA remains unclear. Given this concern, it is perhaps less compelling to use exclusive dsDNA dyes such as CyQuant and picoGreen for AAV titration. On the other hand, GelRed® and GelGreen® dyes bind both ssDNA and dsDNA, making them more suitable than dsDNA dyes for measuring AAV’s genome content.

We have explored two methods for releasing viral DNA from viral particles. The first method is heat inactivation. It has been demonstrated that AAV serotypes are different in their thermal stability, with rAAV2 being the least thermal stable and rAAV5 being the most thermal stable. In PBS buffer, the melting temperature (Tms) of different AAV serotypes range from 66.5 up to 89.5 °C^43^. Therefore we choose to heat rAAV samples at 95 °C. At such high temperature, even the most stable rAAV5 should be destroyed. The second method we used is the alkaline lysis method. In our experiment, we found the EC_50_ of NaOH treatment to be 12 mM for AAV serotype 2. It remains to be examined whether different AAV serotypes share a similar sensitivity to NaOH treatment. To be more rigorous, we have used high concentration of NaOH (100 mM) in our experiments to lyse AAV.

AAV8RSM has been extensively characterized by ARMWG. In the first paper published in 2014^38^, genome titer was determined be 9.62 × 10^11^ GC/ml, based on qPCR data obtained from 16 labs. However, significant variations were found among these labs, with almost 100-fold difference between the lowest titer (4.6 × 10^10^ GC/ml) and the highest titer (4.7 × 10^12^ GC/ml). Elisa method was also used to measure capsid particle titer of AAV8RSM. This assay yielded a value of 5.5 × 10^11^, which is actually lower than the value determined by qPCR^38^. The second paper published in 2016 demonstrated a “free-ITR” qPCR method. Using this method, the titer of AAV8RSM was measured to be 5.65 × 10^11^ GC/ml, which was close to the titer determined by dot blot method and Elisa methods (Table 4, D’Costa et al. ^26^). In the third paper published in 2018, AAV8RSM titer was determined to be 5.65 × 10^11^ GC/ml by qPCR targeting the SV40 polyA sequence (Table 1, François et al.^42^). This result is similar to the results published in 2016^26^. However in 2019, three independent labs from ARMWG carried out AAV8RSM titration again^32^. This time, the genome titer determined by qPCR (1.48 ± 0.618 × 10^12^ GC/m) was 2-3 fold higher than total capsid particle titer determined by Elisa (5.76 ± 0.33 × 10^11^). So despite a series of studies that spanned many years, discrepancies remain to be resolved, although it was evident that ELISA method was more consistent than qPCR method^32^.

We decided to statistically compare the titer determined by GelGreen method in our study to the titers determined by Elisa and qPCR methods in ARMWG’s most recent report^32^. For this purpose, we imported results from Penaud-Budloo et al.^32^ and re-plotted their data as “Elisa” group and “qPCR” group in parallel with our GelGreen data (Figure 7). One-way ANOVA revealed significant difference among the three groups (F_2,7_= 6.053, p<0.05). Tukey’s post-test showed that the GelGreen group is significantly different from the qPCR group but is not different from the Elisa group. Specifically, titer measured by GelGreen was 4.23 ± 0.70 × 10^11^, which is slightly lower although is still within one standard deviation to the titer measured by capsid ELISA (5.73 ± 2.62 × 10^11^).

**Figure 7.**
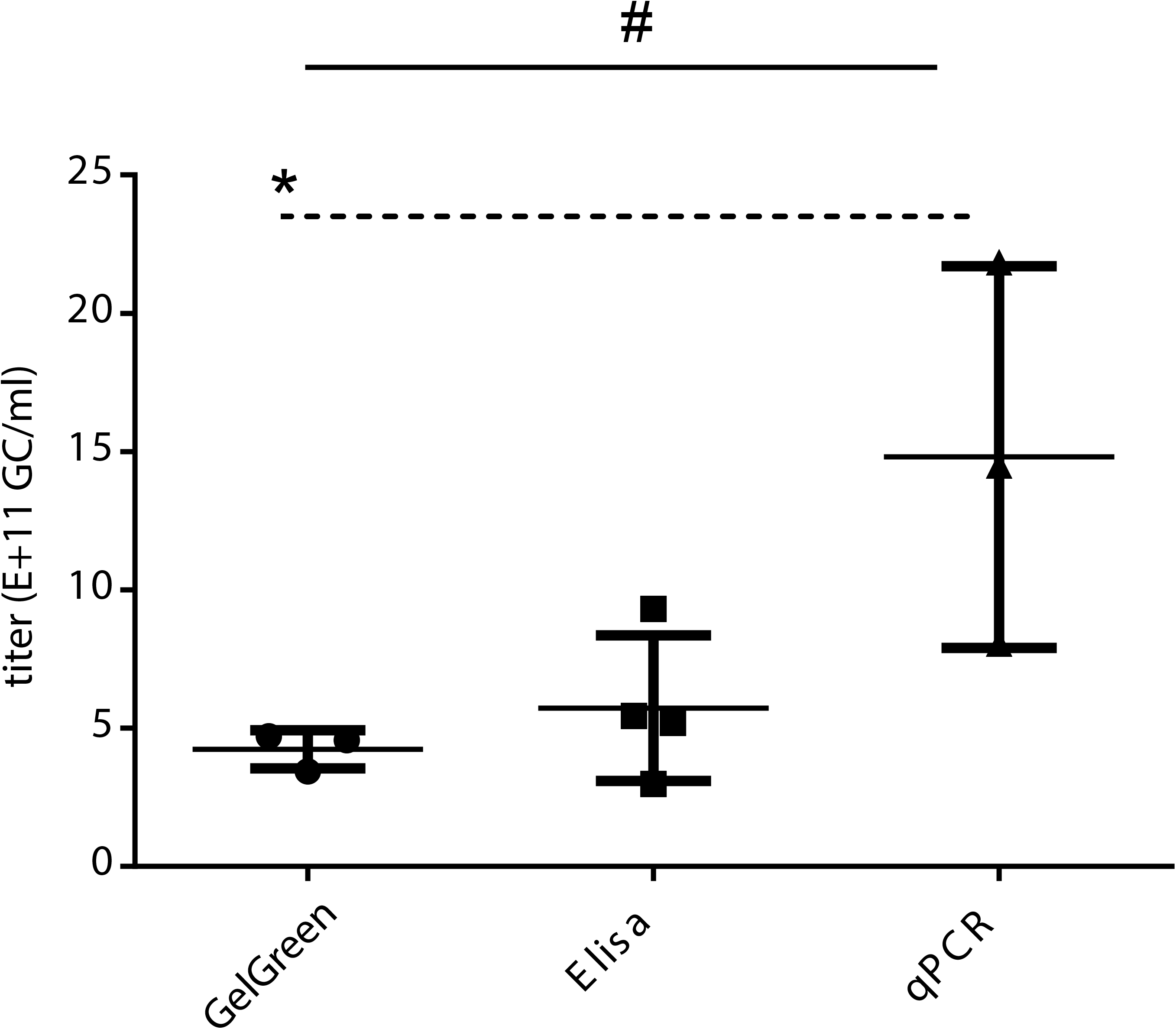
Comparisons of titers of AAV8RSM determined by different methods. Bar graph showing titers of AAV8RSM determined by three different methods. The first set of data is based on GelGreen method in this study. The second set (Elisa) and third set (qPCR) of data was imported directly from Table 3 and Table 1 of Penaud-Budloo 2019^32^. Bars indicate SD. #, p < 0.05, one way ANOVA; *, p < 0.05, Tukey multi-comparison test.

In conclusion, we report a protocol to measure AAV titer using safe nucleic acid dyes. This protocol is simple, safe, reliable and cost-efficient. It could be broadly applied for quantification and normalization of AAV vectors. Future studies can explore more DNA dyes and may find improvements in detection limit.

## Acknowledgements

The authors thank Northwestern University Analytical BioNanotechnology Equipment Core facility of the Simpson Querrey Institute for providing Cytation 3 plate readers and for technical support.

## Author Contributions & Funding Sources

JX, YZ and SHD designed experiments, collected and analyzed data, and wrote the manuscript. This work was supported by NIH R01 EY030169 to YZ, Whitehall grant to YZ and NIH R01 EY018204 to SHD.

